# Catalytic domain-lipid interactions negatively regulate RhoA signaling by modulating conformational dynamics

**DOI:** 10.64898/2026.11.10.765649

**Authors:** Chase M. Hutchins, Yan Zuo, Stephanie A. Planque, Jacob Jakubec, Jeffrey A. Frost, Alemayehu A. Gorfe

**Affiliations:** Department of Integrative Biology and Pharmacology, McGovern Medical School, University of Texas Health Science Center at Houston, 6431 Fannin St., Houston, Texas 77030, USA; Molecular and Translational Biology Program & Therapeutics and Pharmacology Program, MD Anderson Cancer Center UTHealth Graduate School of Biomedical Sciences, 6431 Fannin St., Houston, Texas 77030, USA

**Keywords:** RhoA, small GTPase, membrane orientation, membrane organization, cytoskeleton, biophysics, lipids

## Abstract

RhoA belongs to the Ras superfamily of small GTPases, regulates cytoskeletal actin organization and cell motility, and is implicated in cancer metastasis and neurodegenerative diseases. It is predominantly anchored to the cytosolic side of the plasma membrane (PM) via its prenylated C-terminus, but little is known about its PM organization, dynamics, and lipid interactions. In this work, we used extensive atomistic molecular dynamics (MD) simulations and simulations-guided experiments to study the molecular details and functional consequences of RhoA membrane targeting, interactions and dynamics. We found that both GDP- and GTP-bound RhoA adopt a well-defined, stable membrane organization and binding mode irrespective of initial conditions and bilayer lipid composition. RhoA membrane organization is determined by hydrophobic interaction of the prenylated C-terminus with the membrane, and a previously unknown, critical role of electrostatic interactions between cationic residues on the G-domain and N-terminus with anionic lipids in the PM. N-terminal interactions are mediated by a polybasic patch including residues R5, K6, and K7, and substitution of these residues with Ala increased RhoA-mediated F-actin accumulation and effector activation in cells. MD simulations suggest that the triple mutant retains the global membrane organization of wild type RhoA but alters key protein-lipid interactions. Specifically, we observe a loss of cooperative lipid engagement involving the polybasic patch and Switch II residues with amine-bearing lipids, and a rerouting of intra-protein dynamic networks that primes Switch I for effector recognition. These results suggest that the membrane serves not just as a scaffold but also as an allosteric partner of RhoA to fine-tune intra-protein dynamics in a functionally relevant manner.

## Introduction

RhoA (Ras homolog family member A) is a membrane-associated 22 kDa Ras superfamily small GTPase that cycles between active GTP- and inactive GDP-bound states to mediate cellular processes controlling cell growth, cytoskeletal actin organization, and trafficking^1–4^. In these proteins, GTPase cycling is regulated by a complex framework of proteins consisting of guanine nucleotide exchange factors (GEFs) and GTPase activating proteins (GAPs). GEFs facilitate GDP dissociation while GAPs catalyze GTP hydrolysis to GDP^5,6^. Disruption of GDP/GTP cycling can lead to cancer^7,8^ as well as neurodegenerative^9,10^ and metabolic diseases^3^, with RhoA dysfunction driving metastasis such as in diffuse gastric cancer^11^ and progression of neurodegeneration^12^.

Small GTPases share a conserved catalytic G-domain structure that harbors the nucleotide- and effector-binding regions. As a result, they display a similar biochemistry such as effector and regulator binding via the flexible Switch I and II regions that change conformation upon GDP/GTP exchange^13^. In the Ras and Rho family small GTPases, the G-domain is anchored to the plasma membrane (PM) via lipid modification(s) at the end of an intrinsically disordered C-terminus called the hypervariable region (HVR)^14^. The catalytic domain of RhoA, a founding member of the Rho family, shares 31% sequence identity with KRAS, a prominent member of the Ras family. Additionally, while RhoA’s HVR contains a five-residue polybasic segment similar to KRAS, it is geranylgeranylated^15^ rather than being farnesylated. Moreover, as in other Rho family members, RhoA is distinguished from Ras by a conserved helical insert near the nucleotide-binding site^16^. Once targeted to the PM via a geranylgeranylated HVR, RhoA serves as a signaling hub to regulate diverse cellular processes^17^ including cytoskeletal actin organization and cell motility^18,19^, cell growth and proliferation, and survival^20,21^. Upon activation by growth factors^22^, mechanical stress^23^, or reactive oxygen species^24^, RhoA transduces signals through a number of effector proteins, including Rho-associated coiled-coil kinase (ROCK)^25^ and mammalian homolog of Drosophila diaphanous (mDia)^26^. ROCK stimulates LIM kinase and suppresses myosin light chain phosphatase (MLCP) activity, thereby inhibiting cofilin and promoting actomyosin contraction, respectively^27,28^. RhoA activation of mDia promotes actin polymerization, resulting in long, straight actin fibers^29^. RhoA activity is also controlled by guanine nucleotide dissociation inhibitors (GDI) that sequester the prenyl anchor and thereby prevent membrane binding^30^.

In this work, we combined extensive (80 μs aggregate time) atomistic molecular dynamics (MD) simulations with mutagenesis and functional assays to map the PM interaction and dynamic landscape of RhoA. We found that RhoA is less dynamic than Ras proteins and engages the membrane in a single, stable configuration that is stabilized by electrostatic interactions, including through basic residues at the N- and C-termini. Mutation of lipid-interacting N-terminal cationic residues to Ala resulted in enhanced F-actin accumulation and MLC2 phosphorylation through a membrane-induced modulation of dynamic coupling within the catalytic domain. Our results suggest that catalytic domain-lipid interactions negatively regulate RhoA signaling by modulating the conformational dynamics of its effector binding regions.

## Methods

### Molecular dynamics (MD) simulations

#### Initial structure and system construction

Crystal structures of the wild type GDP-(PDB ID 1FTN) and GSP-bound (1A2B) human RhoA catalytic domain (residues 1-177) were downloaded from the Protein Data Bank (PDB)^31^. The non-hydrolysable GTP-analogue GSP was converted to GTP, missing N- and C-terminal residues were modeled as extended chains, and the C-terminal Cys was modified by geranylgeranylation and carboxymethylation using CHARMM-GUI’s PDBreader^32^. We refer to these two systems as RhoA-GDP and RhoA-GTP-1, respectively. Another initial structure of RhoA-GTP (RhoA-GTP-2) was constructed by rotating the HVR about the membrane normal by ∼60°, followed by a brief local energy-minimization using UCSF ChimeraX. Two RhoA-GTP mutant systems were also built: a triple mutant (R5A/K6A/K7A, hereafter RhoA-GTP-3A) and a double mutant (R5A/K6A, RhoA-GTP-2A). In parallel, we used the CHARMM-GUI Membrane Builder^33^ to prepare two bilayer models that differ in compositional complexity: a simpler model of the PM comprised of 190 POPC lipids in one leaflet and 98 POPC, 60 POPE and 40 POPS lipids in the other to facilitate comparison with previous simulations of KRAS^34^, and another more complex PM model made up of 100 palmitoyl-sphingomyelin (PSM) and 102 POPC lipids in one leaflet and 50 each of POPC, POPE, POPS, and plasmalogen PE (PlaPE) lipids in the other (Table 1). The different numbers of lipids per leaflet in each system were designed to minimize monolayer area asymmetry, as shown in SI, to prevent curvature^35^ and determined based on area per lipid (APL) estimates available in the CHARMM-GUI Membrane Builder. The prenyl chain of RhoA-GDP and RhoA-GTP-1 was then inserted into the mixed-lipid (“lower”) leaflet of the simpler PM model bilayer while RhoA-GTP-2, RhoA-GTP-3A and RhoA-GTP-2A were attached to the more complex bilayer model, as described previously^35^. Each system was solvated with TIP3P waters^36^ and neutralized by Na^+^ and Cl^-^ions for a final concentration of 150 mM, resulting in the five systems listed in Table 1 and illustrated in Fig 1A.

**Figure 1:**
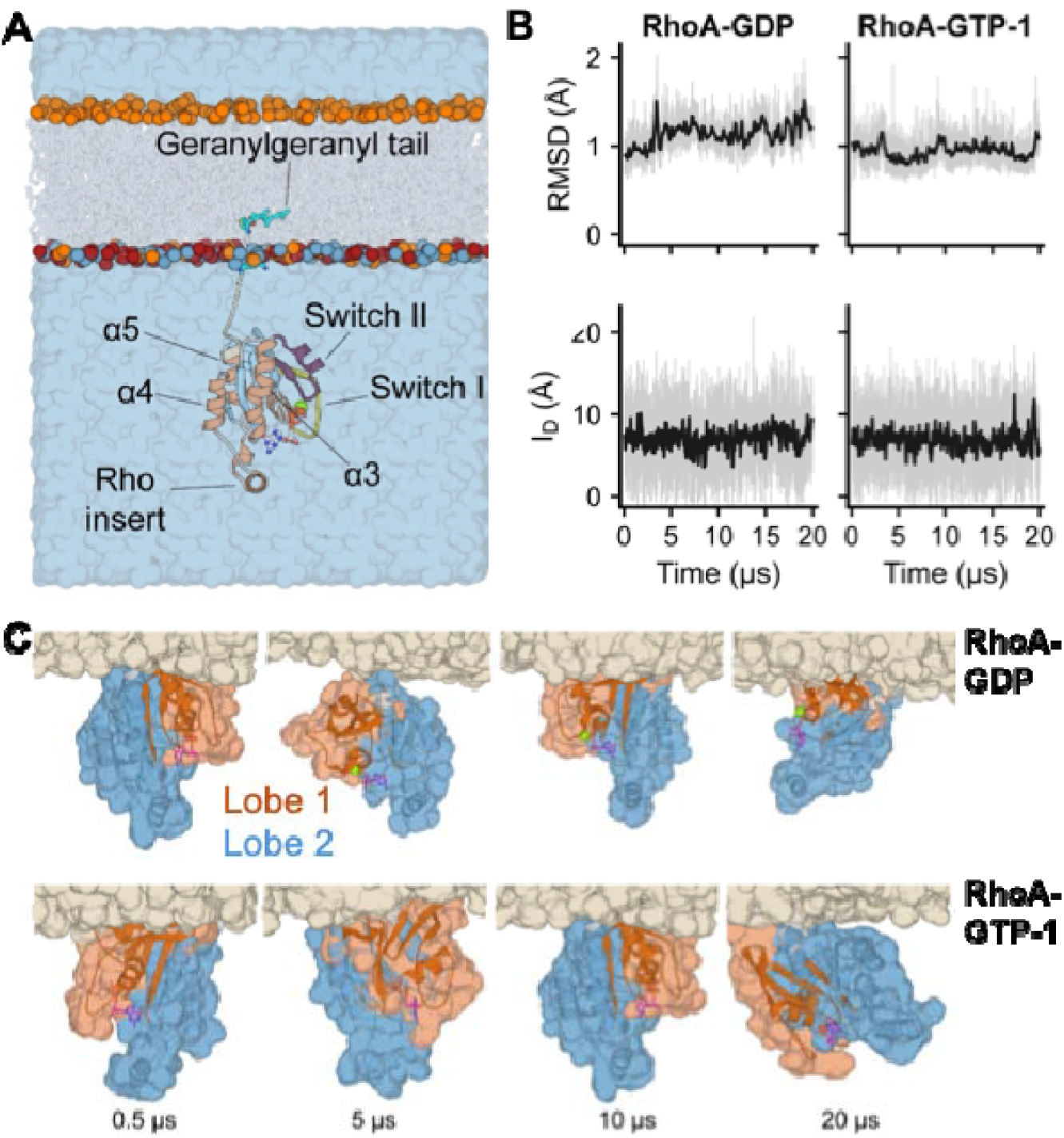
Simulation setup and global dynamics of GDP- and GTP-RhoA in a PC-PC/PE/PS bilayer. **(A)** The RhoA-GDP simulation system with helices in light orange and β-strands in light blue, GDP in stick, Mg²⁺ in green sphere, and geranylgeranyl in cyan. Key structural regions are labeled, including Switch I (yellow; residues 28-42), Switch II (purple; residues 56-78), helices α3 (residues 86-103), α4 (residues 138-148), α5 (residues 164-177), and the Rho insert (residues 122-131). The bilayer is depicted by phosphate atoms to denote lipid identity, with POPC, POPE, and POPS shown as slate, green, and red spheres, respectively, and lipid acyl chains rendered as semi-transparent white sticks. Solvent is shown as a blue shade. (**B**) Time-evolution of catalytic domain (residues 1-180) Cα RMSD excluding the intrinsically disordered switch I and II regions (top) and insertion depth of the geranylgeranyl lipid tail defined as center-of-mass z-distance from the bilayer phosphate plane (bottom). (C) Snapshots at the indicated times illustrating dynamics of RhoA-GDP and RhoA-GTP-1. Lobe 1 (residues 1-86) and lobe 2 (residues 87-177) are in blue and vermillion, a portion of the membrane is shown in sand-colored surface; the hypervariable region (HVR) in light gray is buried in the bilayer. Note the similar fluctuations in the two systems with persistent membrane contacts of the G-domain via lobe 1.

**Table 1.**
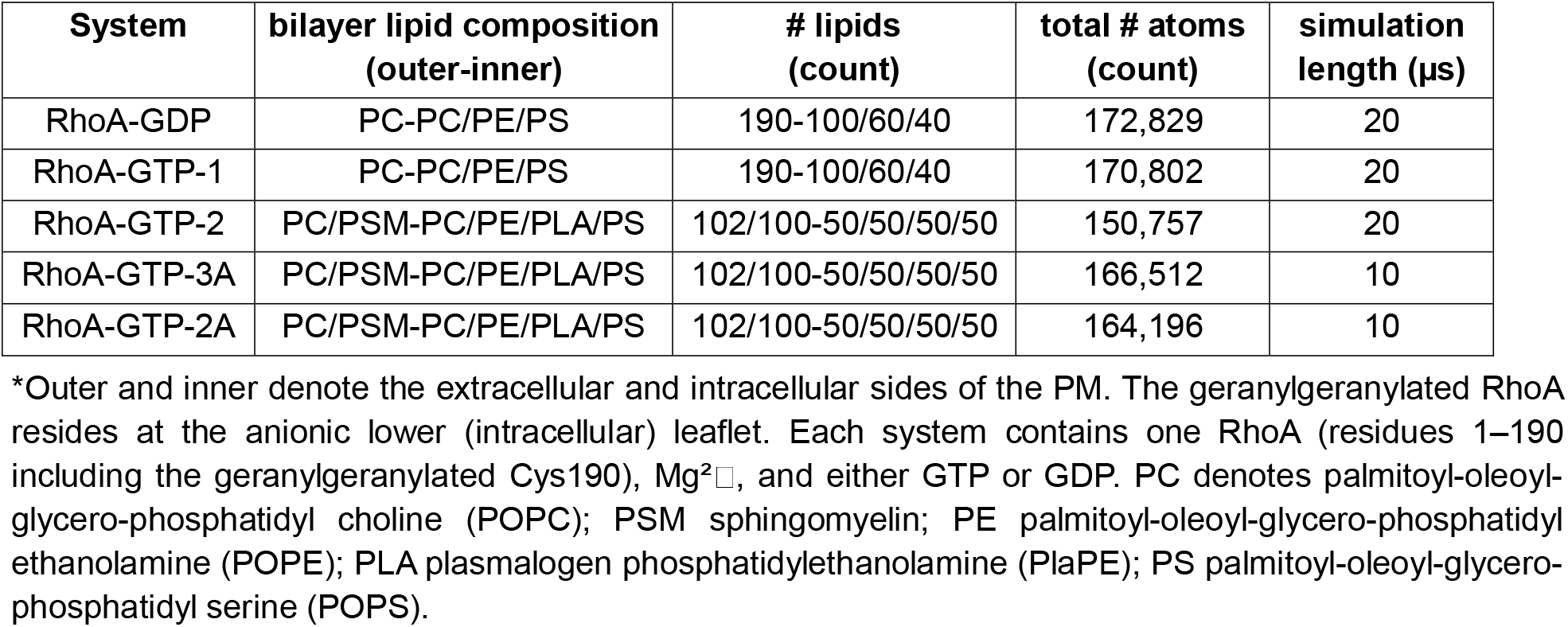
Summary of simulation systems*.

| **System** | **bilayer lipid composition<br>(outer-inner)** | **# lipids<br>(count)** | **total # atoms<br>(count)** | **simulation<br>length (µs)** |
|---|---|---|---|---|
| RhoA-GDP | PC-PC/PE/PS | 190-100/60/40 | 172,829 | 20 |
| RhoA-GTP-1 | PC-PC/PE/PS | 190-100/60/40 | 170,802 | 20 |
| RhoA-GTP-2 | PC/PSM-PC/PE/PLA/PS | 102/100-50/50/50/50 | 150,757 | 20 |
| RhoA-GTP-3A | PC/PSM-PC/PE/PLA/PS | 102/100-50/50/50/50 | 166,512 | 10 |
| RhoA-GTP-2A | PC/PSM-PC/PE/PLA/PS | 102/100-50/50/50/50 | 164,196 | 10 |

\*Outer and inner denote the extracellular and intracellular sides of the PM. The geranylgeranylated RhoA resides at the anionic lower (intracellular) leaflet. Each system contains one RhoA (residues 1–190 including the geranylgeranylated Cys190), Mg²□, and either GTP or GDP. PC denotes palmitoyl-oleoyl-glycero-phosphatidyl choline (POPC); PSM sphingomyelin; PE palmitoyl-oleoyl-glycero-phosphatidyl ethanolamine (POPE); PLA plasmalogen phosphatidylethanolamine (PlaPE); PS palmitoyl-oleoyl-glycero-phosphatidyl serine (POPS).

#### Simulation and trajectory analysis

Using the CHARMM36 force field^37^ and NAMD^38^, each system was energy-minimized using the conjugate gradient method, first for 2000 steps with lipid phosphorus and protein backbone atoms fixed and then for 2000 steps without fixed atoms. This was followed by equilibration with Δt = 1 fs for 2 ns applying a harmonic restraint of force constant k = 4 kcal/mol/Å2 to lipid phosphorus and protein backbone atoms, followed by three steps of 1 ns runs with k reduced by 1 kcal/mol/Å^2^, and an additional 2 ns with k = 0. We then conducted 40-250 ns equilibration runs with Δt = 2 fs using a switching function, with 10 and 12 Å distance cutoffs for non-bonded interactions and a 14 Å cutoff for pair-list update. The NPT (constant number of particles, pressure, and temperature) ensemble was used with the Nosé-Hoover Langevin piston and Langevin thermostat to maintain pressure at 1 bar and temperature at 310 K, respectively. Production runs were conducted on Anton 2 or Anton 3^39^ with default parameters for the NPT ensemble and a 2.5 fs timestep for 20 µs (RhoA-GDP, RhoA-GTP-1 and RhoA-GTP-2) or 10 µs (RhoA-GTP-3A and RhoA-GTP-2A) for an aggregate time of 80 µs. Structures were written out every 100 ps for analysis.

Trajectories were analyzed using MDAnalysis^40^, GROMACS tools, and VMD scripts^41^ while statistical analyses were made using SciPy^42^ on the equilibrated portion of each trajectory. We used Matplotlib^43^ for data visualization and PyMOL^44^ for structure rendering. Standard techniques, such as those described in our recent report^45^ and summarized in SI, were used for analyses of common membrane and protein structural properties. These include bilayer thickness (P-P) and area per lipid (APL); root mean square deviation (RMSD) and fluctuation (RMSF); interatomic distances and contacts; hydrogen bonding (HB) and vdW interactions; membrane orientation angles and distances. Allosteric couplings were probed with the generalized-correlation dynamical network framework^46^ and the Dynetan library^47^, wherein nodes represent residues and coupled motions represent Kraskov–Stögbauer–Grassberger mutual information^48^. Correlations and networks were computed over the last 5 µs data sampled every 0.5 ns, and using a heavy atom distance cutoff of 4.5 Å and contact frequency cutoff of 75%. Communities of tightly coupled residues were detected by Louvain modularity maximization^49^. Other equilibrium membrane and protein properties were computed over the last 15 µs of trajectories. See SI for additional details.

### Mutagenesis and cell assays

#### Plasmids

Full-length coding sequences of HA- and GFP-tagged RhoA were synthesized (Twist Bioscience) and cloned into a pcDNA3.1(-)-Puromycin backbone. Coding sequences and plasmid vector were digested with ApaI and EcoRI restriction enzymes, gel-purified, and ligated with Hi-T4 DNA ligase (NEB). The resulting constructs were transformed into *E. coli* (NEB5α) under ampicillin selection, and the DNA sequences were confirmed with Sanger sequencing.

#### Cell culture

NIH3T3 cells were grown in high-glucose DMEM with 10% fetal bovine serum (FBS) and 1% penicillin/streptomycin (Cytiva, Corning, Gibco) on coverslips in 6-well plates (Corning, CLS3516). Cells were transfected in serum-free medium by adding 100 µL of a 2 μg plasmid and 2 µL Jet Optimus (Polyplus) solution, incubated for 4 h, followed by replacing the medium with complete medium and allowing expression for 24 h before experiments.

#### Immunofluorescent staining and imaging

Cells were grown with serum (10% FBS in DMEM) or serum-starved (overnight in DMEM with 0.5% FBS) prior to fixing. Cells were fixed in 4% paraformaldehyde for 5 minutes at room temperature and washed three times for 5 minutes each with PBS (phosphate-buffered saline: 137 mM NaCl, 2.7 mM KCl, 10 mM Na₂HPO₄, 1.8 mM KH₂PO₄, pH 7.4). Cells were permeabilized with 0.2% Triton X-100 in PBS for 5 minutes, washed three times with PBS, and blocked in 2% goat serum in PBS for 1 hour at room temperature. Coverslips were then incubated overnight at 4°C with mouse anti-HA monoclonal antibody (Cell Signaling Technology, #3724; 1:1000 dilution) and rabbit anti-phospho-myosin light chain 2 (Cell Signaling Technology, #3675; 1:400 dilution) diluted in blocking buffer. After washing with PBS + 0.05% Tween-20, cells were incubated for 1 hr at room temperature with Alexa Fluor 488-conjugated anti-rabbit secondary antibody (ThermoFisher, 1:1000) and Alexa Fluor 555-conjugated anti-mouse secondary antibody (ThermoFisher, 1:1000), together with Alexa Fluor 647-conjugated phalloidin (Cytoskeleton Inc., 1:500) for F-actin visualization. After three final PBS+ 0.05% Tween-20 washes, coverslips were mounted onto slides using Fluorsave mounting medium (ThermoFisher).

Images were acquired on a Zeiss Axiophot epifluorescence microscope using a 60× 1.4 NA oil-immersion objective and Axiovision acquisition software. Cells with no detectable HA signal were excluded from analysis. Images were analyzed in Fiji/ImageJ with regions of interest (ROIs) drawn manually around individual transfected cells using the HA channel as a guide. Background was subtracted from each channel using the rolling-ball algorithm prior to measuring mean fluorescence intensity within each ROI. For each cell, phalloidin or pMLC2 mean intensity was divided by HA mean intensity within the same ROI to yield an expression-normalized signal per cell. Normalized intensities from independent biological replicates were pooled for statistical analysis in GraphPad Prism using one-way ANOVA with Tukey post-hoc comparisons.

## Results

Mammalian cell PM is compositionally asymmetric with its cytosolic side, where RhoA predominantly resides, enriched with PE and PS lipids^50^. We therefore conducted atomistic MD simulations of GDP- and GTP-bound RhoA in asymmetric 3-component PC/PE/PS and 5-component PC/PSM/PE/PLA/PS bilayers, detailed in Table 1 and Fig 1A (see Methods). We compared the results with previous simulations of RhoA, KRAS and Rheb, which allowed us to assess whether RhoA membrane engagement and dynamics are primarily encoded in the primary sequence or are a consequence of host membrane complexity, nucleotide state, or initial conditions. In subsequent sections, we will first present results from the simulations in the 3-component bilayer and then the 5-component bilayer, followed by data from experiments and mutant RhoA simulations.

### RhoA is characterized by restricted orientational dynamics in a 3-component lipid bilayer

Fig. 1B displays the time evolution of Cα RMSD (excluding the flexible switch regions and the chain termini) and the insertion depth of the geranylgeranyl acyl chain (l_D_) in simulations RhoA-GDP and RhoA-GTP-1. The data demonstrate equilibration of both systems within ∼5 μs, as indicated by the plateauing of the plots at RMSD 1.2 Å and l_D_ ≈ 7.5 Å past the phosphate plane of the mixed-lipid monolayer. The bilayer equilibrated even faster as assessed by APL and P-P measurements (Fig. S1), consistent with previous observations^51^. While the RhoA G-domain remained attached to the bilayer via the geranylgeranyl moiety throughout the simulations, the snapshots in Fig. 1C show local fluctuations while lobe 1 (residues 1-86) remained proximal and lobe 2 distal to the membrane surface. This is qualitatively consistent with a previous observation in a simpler 80/20 PC/PS symmetric bilayer^52^. The apparent lack of global membrane orientational motion in the current simulations is supported by quantitative data based on the COM z-distance of lobe 1 (Zlb1) and lobe 2 (Zlb2) from the bilayer center (Fig. 2). Z_lb1_ fluctuated extensively but remained at an average value of < 40 Å while Z_lb2_ fluctuated less and stabilized at ∼50 Å (Fig. 2A). GDP-bound RhoA is generally more dynamic than its GTP-bound counterpart in terms of both RMSD and Z_lb_ distances (Figs. 1 & 2). However, in both cases Z_lb1_ versus Z_lb2_ 2D distributions based on the equilibrated last 15 μs data show a single dominant peak centered at Z_lb1_ ≅ 40 Å and Z_lb2_ = 48-49 Å (Fig 2B). This shows that, irrespective of the nucleotide state and unlike the RAS family proteins KRAS^34,53^, HRAS^54^ or Rheb^45,52,55^, the G-domain of RhoA does not undergo re-orientation on the 3-component bilayer: lobe 1 (also called effector lobe) is membrane proximal and lobe 2, or the allosteric lobe, is distal despite occasional attempts at re-orientation.

**Figure 2:**
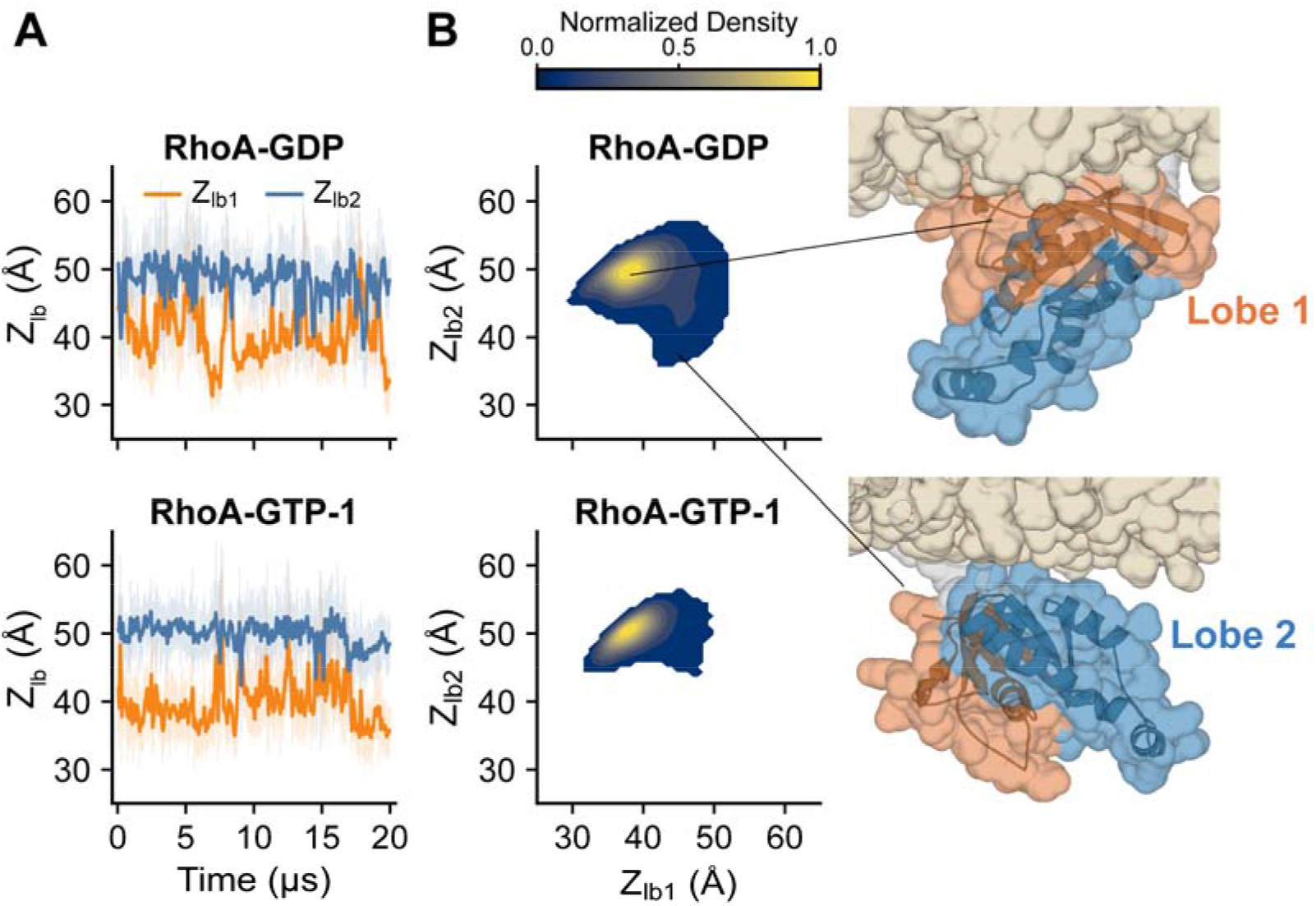
RhoA G-domain dynamics on a bilayer surface. **(A)** Time evolution of the z-distance between the bilayer COM and lobe 1 (Z_lb1_, residues 1-86) and lobe 2 (Z_lb2_, residues 87-177) in blue and gold in RhoA-GDP and RhoA-GTP-1 simulations. **(B)** Normalized 2D density distribution of Z_lb1_ vs. Z_lb2_ for each system. Representative snapshots of membrane-proximal configurations corresponding to the indicated locations on the density map of RhoA-GDP are shown on the right with the bilayer and each lobe colored as in Fig 1.

### Lack of membrane re-orientation is independent of initial condition or lipid composition

To assess the robustness of the observations above and the potential impact of membrane lipid composition or initial condition on RhoA dynamics, we conducted an independent simulation of GTP-bound RhoA in a more complex bilayer (system RhoA-GTP-2 in Table 1). This simulation was started from a different initial configuration, where the HVR of RhoA-GTP-1 was rotated by 60° so that lobe 2 faces the bilayer before the protein was attached to the bilayer (see Methods). As shown in Fig. S1, this 5-component bilayer is characterized by a slightly larger average P-P than the 3-component bilayer (∼40.0 vs. 38.3 Å), as expected by the condensing effect of PSM. The APLs are correspondingly smaller (e.g., 64-66 vs. 61-63 Å^2^ for the upper and lower leaflet POPC lipids). Despite these differences in bilayer properties, RMSD and l_D_ of RhoA in this simulation behaved nearly identically to that in RhoA-GTP-1 (Fig. 2; Fig. 3 A,B). The Z_lb_ distances underwent dramatic changes at ∼5 μs, whereby the initially membrane proximal lobe 2 moved away and lobe 1 approached the membrane surface (Fig. 3C). After this transition, Z_lb1_ and Z_lb2_ remained fluctuating around the equilibrium distances observed in the 3-component bilayer systems of Fig. 2, with no sign of a reverse transition (Fig. 3C). The *p*(Z_lb1,_Z_lb2_) plots encompassing the last 15 μs data also showed a similar profile to that in the RhoA-GTP-1 system, albeit with a somewhat narrower distribution centered at (Z_lb1_, Z_lb2_) = (38,48). The slightly smaller Z_lb1_ value in this simulation suggests a closer lobe 1-bilayer contact, consistent with the somewhat larger protein-membrane contact surface formed following the conformational transition (Fig. 3D). Despite these minor differences, the similar *p*(Z_lb1,_Z_lb2_) equilibrium distributions in all three simulations support the conclusion that RhoA adopts a single membrane organization and orientation at equilibrium, wherein the effector binding regions are positioned adjacent to the membrane surface (Fig. 1C, Fig. 2B, Fig. 3E). Occlusion of the effector binding surface by the membrane has been observed previously in some orientation states of KRAS^56,57^ and Rheb^45,55^. It is important to emphasize that absence of global orientational dynamics does not mean the absence of fluctuations within the protein or in protein-lipid interactions. This is the case in the simulations, as exemplified by the weakly anti-correlated distance fluctuations of Switch I and α4, which are located at opposite faces of the G-domain, from the bilayer surface (Fig. S2). These observations strongly suggest that RhoA engages lipids using a single, well-defined surface patch with its G-domain undergoing small amplitude rocking motions about the membrane interface within a fixed overall configuration.

**Figure 3:**
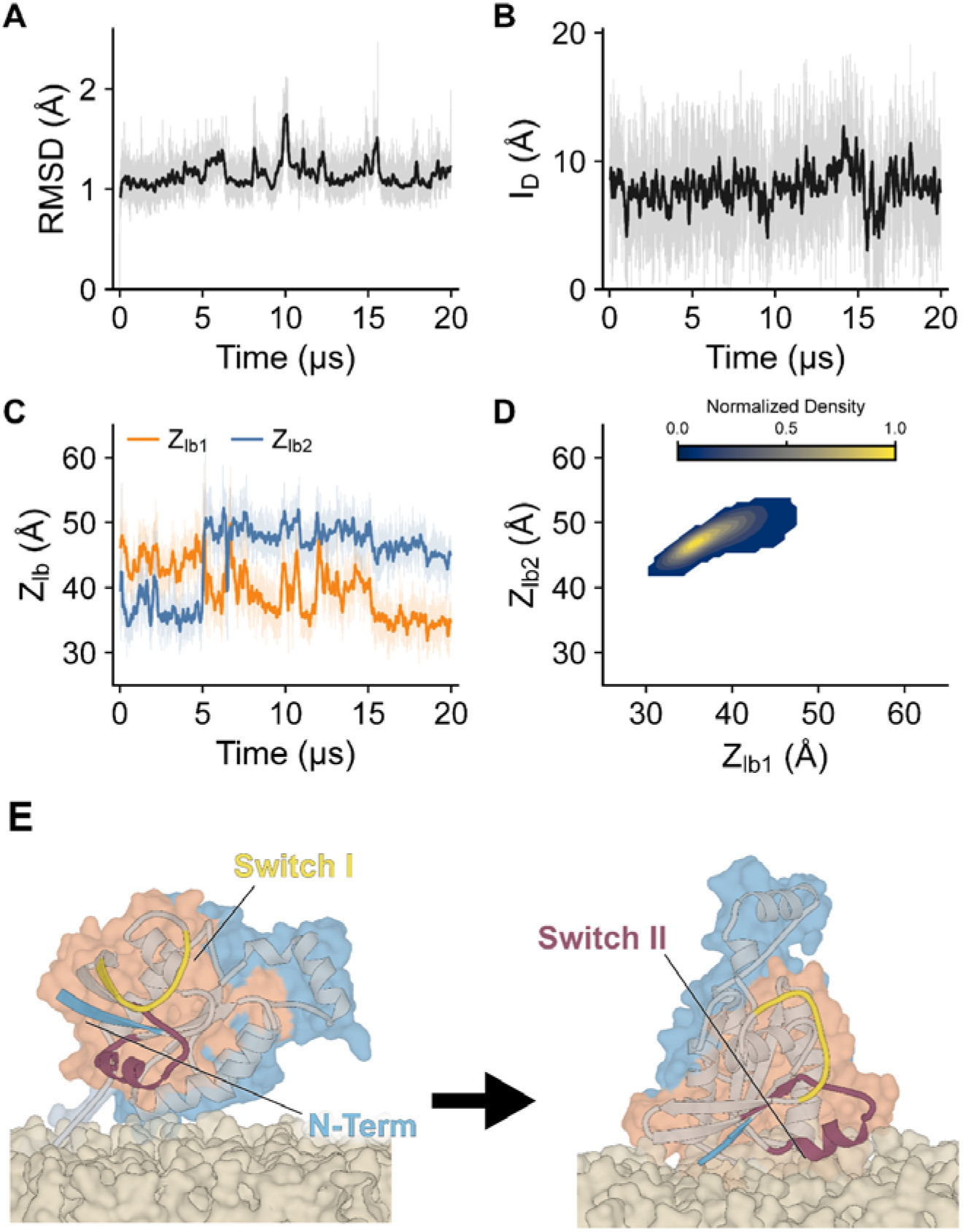
Dynamics and G-domain reorientation during simulation RhoA-GTP-2. (A, B) Time-evolution of the catalytic domain Cα RMSD excluding switch I and II (A) and insertion depth of the geranylgeranyl lipid tail (**B**). (C) Time-evolution of Zlb1 (orange) and Zlb2 (blue). **(D)** Normalized 2D density distribution of Z_lb1_ vs. Z_lb2_. (**E**) The starting (left) and final (right) configuration of RhoA-GTP-2 illustrating the reorientation of the G-domain. Color codes as in previous figures.

### RhoA membrane binding is characterized by specific tilt and rotation angles and involves lipid-interaction of the N-terminus and switch loops

To more quantitatively describe membrane dynamics, we measured tilt (θ) and rotation (φ) angles that describe the inclination of the G-domain from the membrane normal and its azimuthal orientation, respectively (Fig. 4A; see SI). Consistent with the Z_lb_ distances (Figs. 2A, 3A), the time series of θ and φ (Fig. 4B) and the resulting *p*(θ, φ) density maps (Fig. 4C) indicate fluctuations around specific angles but no large-scale transition throughout RhoA-GDP and RhoA-GTP-1 and after 5 μs of RhoA-GTP-2 (i.e., following the conformational transition noted above). This shows not only that the distance- and angle-based reaction coordinates describe the same essential dynamics but also the tendency of the RhoA G-domain to engage lipids in a specific configuration. This configuration is defined by an average θ of 46.5 (± 14)° and φ ≈ -8 to -38 (± 35-44)° in all three systems, indicating a similar global orientation with minor effects of nucleotide state or lipid composition. Analysis of residue-lipid contact frequencies suggests that this orientation is associated with persistent lipid interactions of a few residues at lobe 1 (Fig. 4D). Only 4-8 residues have contact frequency ≥ 0.5 and about a dozen ≥ 0.3, including a cluster of non-polar and basic amino acids at the N-terminus (A_2_A_3_I_4_R_5_K_6_K_7_) and, to a lesser extent, at switches I and II (K51, D76 and D78). As might be expected purely from electrostatics, R5, K6 and K7 are engaged in interactions with the anionic PS lipids (Fig. 4 D,E). These results confirm that RhoA adopts a stable membrane configuration independent of nucleotide identity, starting configuration, or membrane lipid composition due largely to electrostatic interactions of the G-domain with anionic lipids.

**Figure 4:**
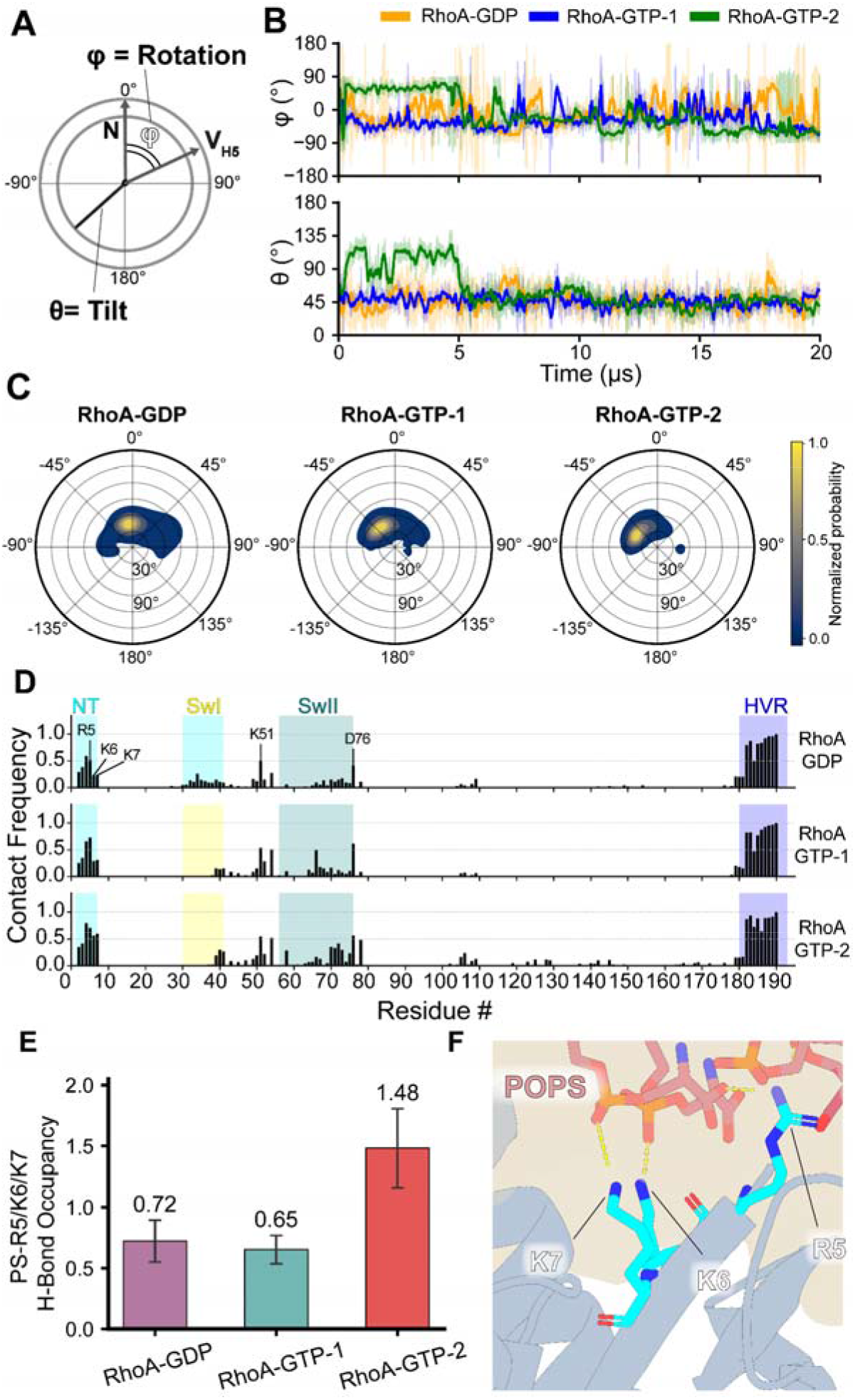
RhoA adopts a single orientation irrespective of initial condition, nucleotide state or membrane lipid composition. **(A)** Definition of membrane-referenced orientation coordinates used throughout this study: rotation angle (φ) defined as the azimuthal angle about the membrane normal (N) and tilt angle (θ) defined as the polar angle between V_H5_ and the membrane normal (see SI for details). **(B)** Time-evolution of φ (top) and θ (bottom) in the three simulations. In RhoA-GTP-2, the angles underwent transition at 5 μs (red dashed line) and converged to values in the other simulations. **(C)** Probability densities of φ vs. θ with grid lines shown every 45° in φ and 30° in θ. Densities are normalized independently for each simulation. **(D)** Frequency of residue-lipid contacts defined as the fraction of trajectory frames in which any side chain heavy atom is within 4 Å of a lipid heavy atom. NT = N-terminus; SwI = Switch I; SwII = Switch II; HVR = hypervariable region. (**E**) The combined ensemble-averaged number of residue-PS hydrogen bonds for R5, K6, and K7. **(F)** An example of hydrogen bonding interaction between the N-terminal basic residues and PS lipids.

### The functional role of the N-terminal cationic patch: a negative regulator of signaling?

The simulations predicted a RhoA G-domain orientation in which the effector binding surface is occluded by the membrane. If electrostatic interactions of R5, K6 and K7 with lipids are a significant contributor to this ‘occluded’ configuration, then their removal through mutation would result in the relaxation of RhoA-bilayer engagement or altered internal dynamics. One possible manifestation of such an effect is a change in RhoA signaling function. To test this hypothesis, we generated N-terminally GFP- and HA-tagged triple alanine mutants of R5, K6, and K7 in both wild type and constitutively active G14V backgrounds (RhoA-WT-3A and RhoA-G14V-3A, respectively). As noted in the *Introduction*, activation of RhoA promotes F-actin assembly through its effectors ROCK and mDia ^58–60^, which promote accumulation of phosphorylated myosin light chain 2 (MLC2) and F-actin^28,61^. We therefore quantified pMLC2 levels and F-actin accumulation as functional readouts of RhoA signaling. In these experiments, we contrasted effects in cells grown in full serum so that all upstream signaling was retained, as well as in cells serum-starved overnight to reduce upstream signaling. Fluorescence microscopy imaging using phalloidin staining showed increased F-actin levels in serum-starved NIH3T3 cells expressing the GFP-tagged mutant constructs under both wild type and G14V backgrounds (Fig. S3). Experiments with the HA constructs yielded similar results (Fig. 5). In the presence of 10% FBS, the mutations significantly increased F-actin levels relative to controls under both wild type and G14V backgrounds, and pMLC2 levels in the wild type background. A similar trend was found under serum-starved conditions, suggesting that the mutation’s effect is likely independent of upstream signaling. We conclude that loss of positive charges at the N-terminus enhances RhoA signaling independent of serum stimulation, suggesting that interactions between the catalytic domain and the membrane act as a negative regulator of effector engagement.

**Figure 5:**
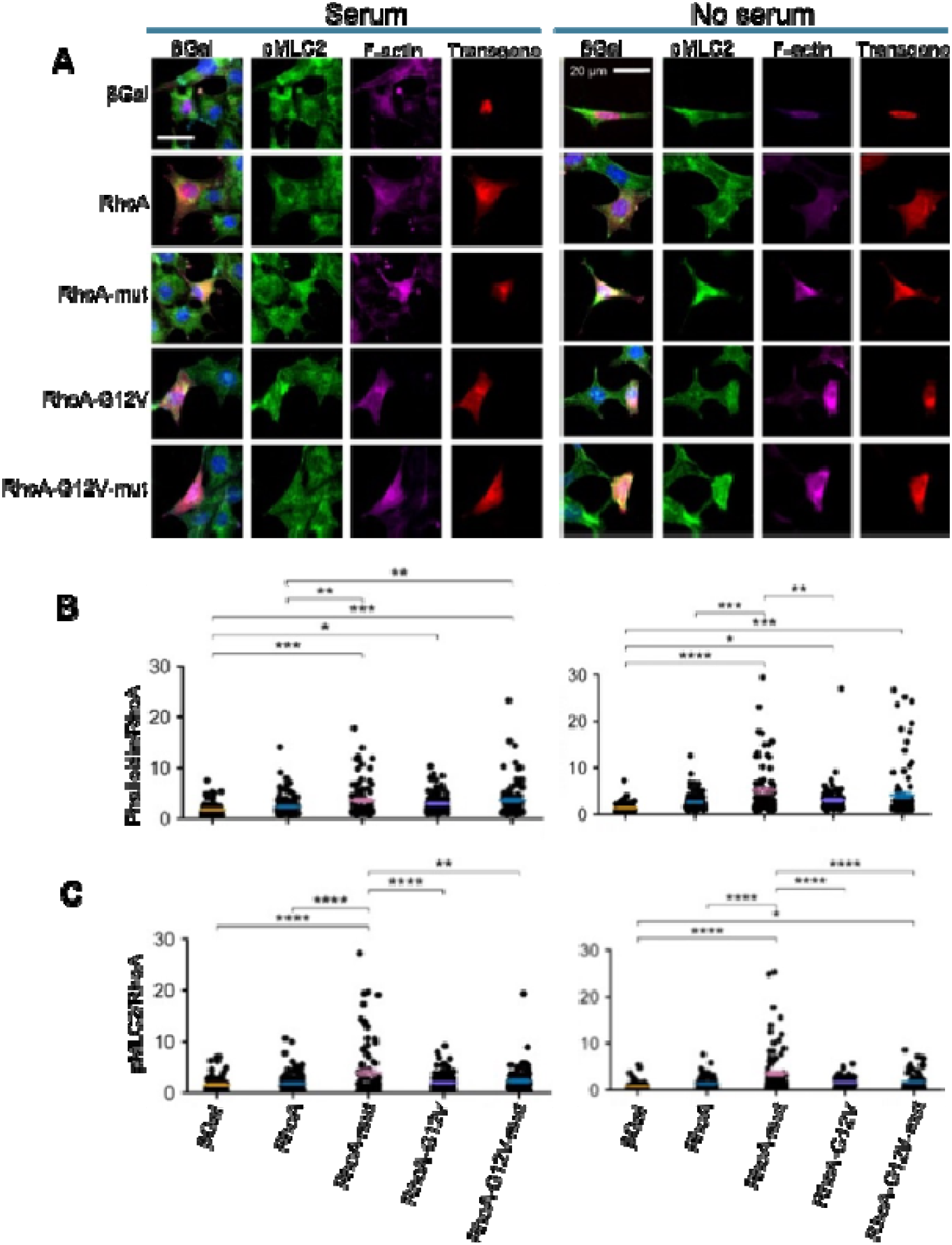
Perturbation of G-domain-lipid interaction modulates RhoA function. **(A)** Representative epifluorescent images of NIH3T3 cells expressing βGal (control), wild type RhoA, RhoA-WT-3A (R5A/K6A/K7A), RhoA-G14V, and RhoA-G14V-3A under serum-fed and no-serum conditions. Cells were stained for βGal or HA-tagged transgene (red), phosphorylated myosin light chain 2 (pMLC2, green), and F-actin (phalloidin, magenta). Scale bar, 20 µm. **(B-C)** Quantification of F-actin **(B)** and pMLC2 (**C**) fluorescence intensities normalized to RhoA expression on a per-cell basis. Each dot represents an individual cell from 4 independent experiments and horizontal bars are median ± SEM. Statistical significance from one-way ANOVA with Tukey post-hoc comparisons: *P < 0.05; **P < 0.01; ***P < 0.001; ****P < 0.0001.

### Ala mutation of the N-terminal polybasic patch remodels lipid interaction and internal dynamics of RhoA

To better understand the enhanced signaling of RhoA-WT-3A (Fig. 5), we conducted a 10 μs MD simulation of the same mutant in the GTP-bound form and in the same 5-component lipid bilayer used in RhoA-GTP-2 (termed RhoA-GTP-3A; Table 1; see SI for a control simulation RhoA-GTP-2A). The mutant remained structurally stable during the simulation and underwent the same conformational changes observed in RhoA-GTP-2, with the initially membrane-distal lobe 1 rotating to face the membrane (Fig. 6A). Following this transition, the global orientational dynamics and membrane engagement of the triple mutant remained very similar to the wild type. This argues against a purely occlusion-based explanation for its increased activity, which prompted us to more closely interrogate protein-lipid interactions and internal dynamics. To this end, we examined the hydrogen bonding of charged residues before, during, and after conformational transition in each system (Fig. 6B). In the wild type reference simulation, R5, K6, and K7 plus switch I and inter-switch residues E40, K51 and E54 are engaged with lipids during and after the orientational transition. On the other hand, all but K51 and E54 interactions are lost in the triple mutant RhoA-GTP-3A post-transition, while interactions of Switch II residues D67, R68, R70 and D76 became more prominent (Fig. 6B). For comparison, the HVR residues engaged lipids in a similar manner in the wild type and mutant RhoA and across conformational states.

**Figure 6:**
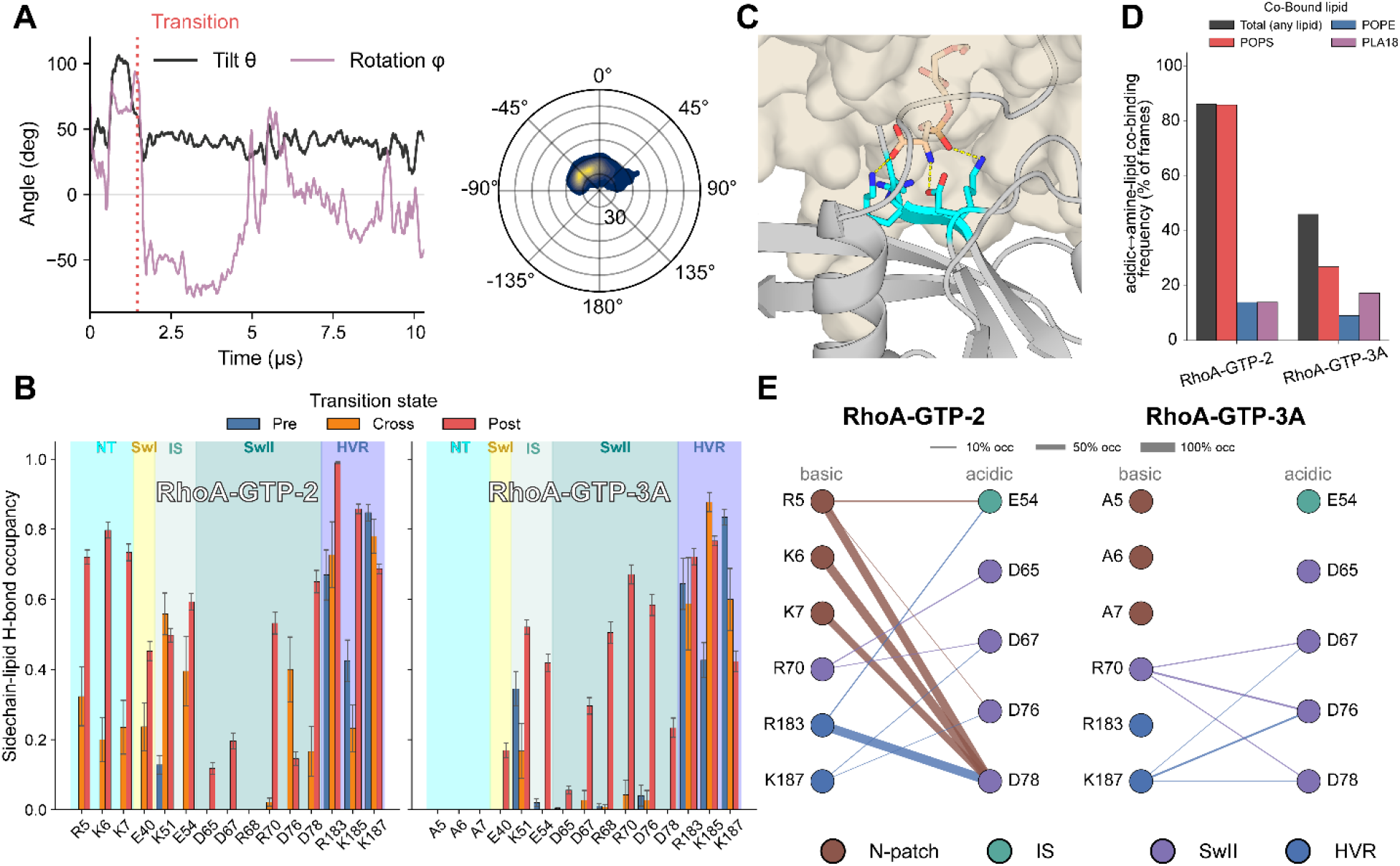
Lipid co-binding of basic and acidic residues. **(A)** Membrane orientation of RhoA-GTP-3A: tilt (θ) and rotation (φ) time series (left; transition into a stable membrane configuration is marked by the dotted line) and the resulting density plot post-transition (right; radial = θ, angular = φ). **(B)** Side chain-lipid hydrogen bond frequency for RhoA-GTP-2 and RhoA-GTP-3A involving residues at the N-terminus, Switch I, inter-switch, Switch II, and HVR regions separately for pre, during and post conformational transition (mean ± SEM). **(C) A snapshot illustrating lipid co-binding** where R5, K6, and K7 (cyan) grip the phosphate of a POPS while D78 grips at the amine. (**D**) Frequency of co-binding averaged over the final 5 µs of RhoA-GTP-2 and RhoA-GTP-3A colored by lipid type. **(E)** Selected residue pairs bridged by a lipid; edge width = co-binding occupancy colored by region. The dense N-patch↔D78 bridging in the wild type collapses in RhoA-GTP-3A.

Since anionic lipids typically engage basic residues^62^, the concomitant increases in Lys/Arg-lipid and Asp/Glu-lipid contacts noted above suggest that the two might be physically proximal and share lipid(s) rather than interacting independently of each other. We tested this with the analysis of co-binding events, defined as a basic and an acidic residue simultaneously interacting with the same lipid molecule (Fig. 6C). We found that, while such events are prevalent in both trajectories, there are nearly twice as many in RhoA-GTP-2 as in RhoA-GTP-3A (86% vs. 46%; Fig 6D). As expected, these events are dominated by PS lipids (only PS, PE, and PlaPE bear a primary amine capable of engaging an acidic side chain). The residue pairs involved in lipid co-binding are shown in Fig. 6E, classified by the strength of the pairing. Each of the mutated residues R5, K6, K7 is strongly (66-86%) coupled with D78 through a shared PS. Not only did the alanine substitutions abolish these lipid-bridged networks, but they also reduced or rearranged other pairs (Fig. 6E). To determine whether this might remodel protein dynamics, we computed generalized-correlation (mutual-information) dynamical networks for each trajectory and partitioned them into communities of concertedly moving residues^47,48^ (Fig. 7A). The N-terminus forms a dynamically isolated node in RhoA-GTP-2 that is condensed into the Interswitch community in RhoA-GTP-3A. The betweenness centrality of D76 and D78 likewise increased in RhoA-GTP-3A (Fig. 7B), identifying them as communication hubs in the rewired network. Consistent with these rearrangements, the N-terminus to Switch I communication path is shorter in the latter, suggesting a tighter coupling between the N-terminus and the effector binding switch loops (Fig. 7C). Since Switch I constitutes the core of the effector binding interface^63,64^, we examined whether the altered lipid-bridged residue networks affect Switch I dynamics as well. Comparison of the simulated Switch I conformers with the crystal structures of RhoA complexes with effectors ROCK1, PKN and anillin indicated major differences between RhoA-GTP-2 and RhoA-GTP-3A (Fig. 7D). That the latter samples conformations closer to the effector-bound Switch I experimental structures is consistent with the mutant’s increased activity in cells.

**Figure 7:**
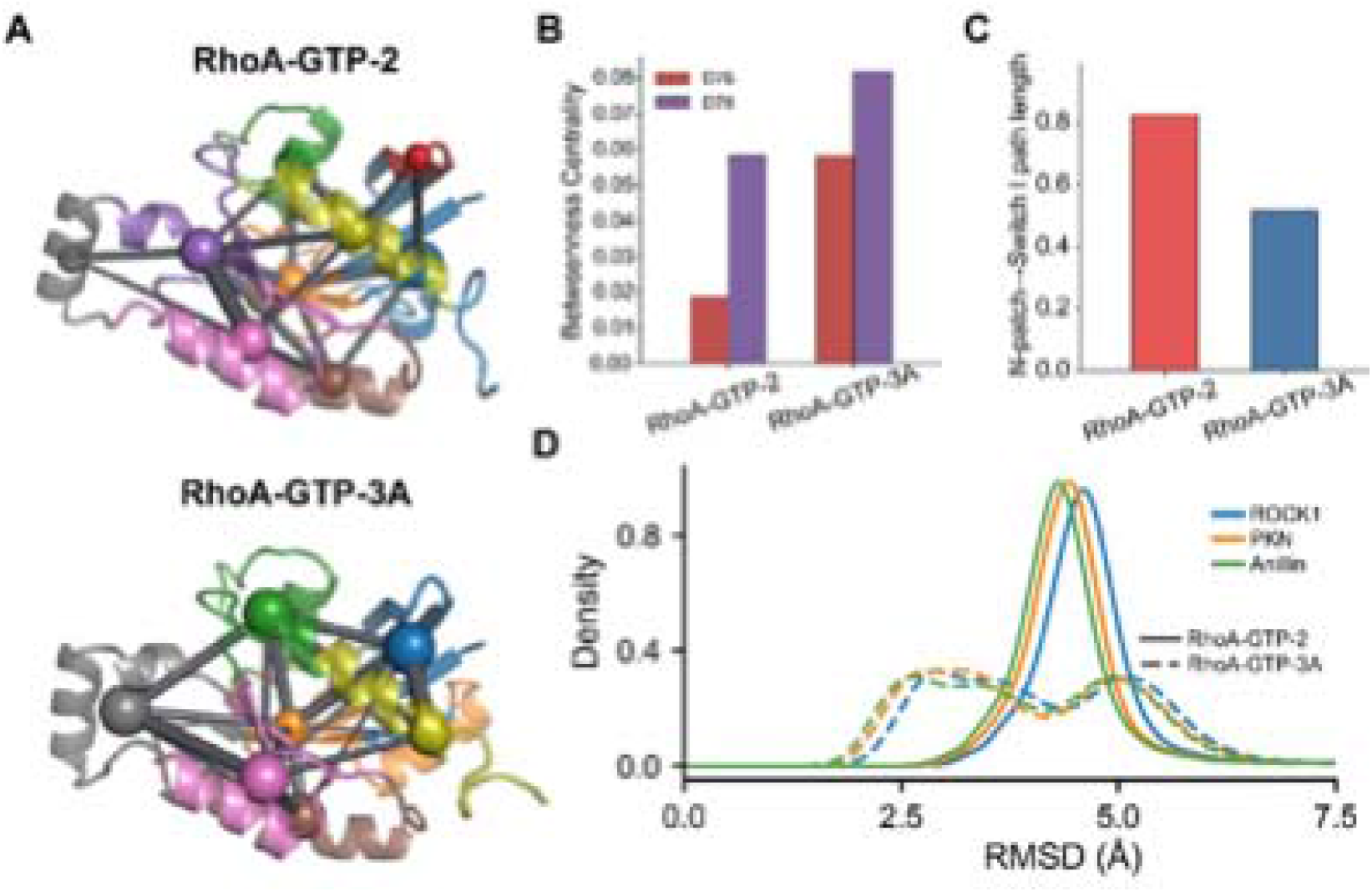
Loss of cooperative lipid co-binding reroutes dynamic couplings and alters Switch I conformational landscape. **(A)** Dynamical communities drawn on the structure for RhoA-GTP-2 (top) and RhoA-GTP-3A (bottom): cartoon colored by community (each residue colored by the wild-type RhoA-GTP-2 community that its own community overlaps with most, so a given module is the same color in both systems), with one sphere per community at its Cα centroid (radius scaled by membership) and rods between communities scaled by their summed inter-community mutual-information coupling. **(B)** Node betweenness centrality of the Switch II residues D76 and D78, which become communication hubs and increase in RhoA-GTP-3A. **(C)** Optimal-path length between the N-terminus and Switch I in the generalized-correlation network (lower = stronger coupling); the channel shortens (strengthens) from RhoA-GTP-2 to RhoA-GTP-3A. **(D)** Switch I Cα RMSD of RhoA-GTP-2 (solid) and RhoA-GTP-3A (dashed) from the effector-bound RhoA conformation of RhoA-effector crystal structures (ROCK1, PDB 1S1C; PKN, 1CXZ; anillin, 4XOI), after superposition on the invariant G-domain core.

## Discussion

In a previous simulation of RhoA in a symmetric PC/PS bilayer, we found that the rotational dynamics of the G-domain were generally less flexible than those observed in the RAS family proteins HRAS^54^, KRAS^34,53^ and Rheb^45,52^. Despite a common protein fold, the Ras and Rho families differ in sequence (e.g., RhoA and KRAS share only 31% sequence identity). Moreover, the previous simulation was conducted in a simplified 2-component model membrane while the PM is compositionally asymmetric, with the cytosolic side enriched in PS and PE lipids^52^. To study whether the unique membrane dynamics of RhoA are encoded in its primary sequence and not a consequence of host membrane complexity, nucleotide state, or initial conditions, we conducted 20 μs atomistic MD simulations each of GDP- and GTP-bound RhoA in an asymmetric PC-PC/PE/PS bilayer and another run of RhoA-GTP in a more complex PC/PSM-PC/PE/PLA/PS bilayer started from a different initial configuration. We found that RhoA adopts a single and stable membrane configuration, independent of initial condition, nucleotide identity, or host membrane complexity. Contact and hydrogen bonding analysis showed that the N-terminal R5, K6, K7 stabilize this configuration through electrostatic interactions with PS lipids and often are engaged in same-lipid interaction with D76 and D78 of Switch II. Alanine substitution of this cationic patch increased F-actin content and effector activation in cells, whereas MD simulation found that the mutant behaved like the wild type in terms of membrane organization. A detailed analysis found rewiring of internal dynamics in the mutant, linking the N-terminus to Switch I through the Interswitch region, thereby altering the ensemble in favor of binding competent conformations.

That the RhoA G-domain adopted a single, highly stable membrane configuration contrasts with the dynamic, multi-state orientation landscapes reported for Ras family proteins^34,53,54,57^. In Ras proteins, orientation dynamics are governed by HVR flexibility and transient interactions between G-domain residues and the membrane^52,53^. In RhoA, the N-terminus acts as a secondary membrane anchor that helps to stabilize a single primary orientation. The conservation of R or K at position 5 across RhoA, RhoB, RhoC, Rac1, Cdc42, Rac2, and RhoG suggests that N-terminal membrane engagement may be a shared feature of the Rho family proteins. Underscoring the importance of this position, R5 of RhoA is a recurrent mutational hotspot in diffuse gastric cancer^65^, where the R5W substitution acts as a gain-of-function mutant that stimulates stress fiber formation and focal adhesion assembly^66–68^. Although this bulky tryptophan substitution likely acts through a partly distinct mechanism, its activating effect is consistent with our finding that perturbing the N-terminal basic patch enhances RhoA signaling. Notably, while C-terminal HVR polybasic regions of Rac1 and Cdc42 have been studied as determinants of membrane localization and signaling^69–71^, functional studies specifically addressing N-terminal basic residues in these proteins or RhoA are lacking. Our observations raise the possibility that N-terminal electrostatic contacts with anionic lipids may be involved in membrane organization across the Rho family, though this remains to be tested directly.

Previous studies of Ras^56,57^ and Rheb^45^ have shown that dynamic membrane occlusion of the effector binding region can impair signaling. In HRAS and KRAS, point mutations that perturb contacts between the catalytic domain and membrane altered signal output and effector engagement^57,72^. Simulations have also shown that orientation preferences in lipid-modified GTPases are determined by the identity and chemistry of residues on the catalytic domain and the HVR^50,52^. The N-terminal basic patch in RhoA may play an analogous role by altering cooperative residue-lipid interactions that result in altered internal coupled dynamics. Moreover, basic residues are well established as key contributors to lipid recognition by peripheral membrane proteins. This includes sequestering PS and PIP2 into laterally enriched pools^62,73^. Acidic residues are rarely described in this role, and thus persistent hydrogen bonds formed by D76 and D78 with amine-bearing lipids were unexpected. These contacts are cooperative rather than independent, with a basic and an acidic side chain hydrogen bonding to the same lipid simultaneously. Such co-binding occurred in 86% of frames in the wild type simulation RhoA-GTP-2 and 46% of RhoA-GTP-3A. This phenomenon is lipid species dependent and thus varies with membrane lipid composition, which parallels the composition-dependent differences in membrane engagement and dynamics reported for KRAS^74^. In the wild type RhoA, R5, K6, and K7 each bridge D78 through a shared PS in 66-86% of frames, which is lost in the mutant and thus modulates the internal dynamics of the G-domain. A similar mechanism of allosteric signals from membrane rerouting intra-protein dynamics has been observed in other small GTPases such as Arf1, where membrane insertion of the N-terminal amphipathic helix is accompanied by a rearrangement of backbone hydrogen bonds in the G-domain core^75^. This is also consistent with the increasingly recognized role of lipids as allosteric effectors rather than passive solvent, promoting receptor-G protein coupling in class A GPCRs^76^, shifting the conformational equilibria of membrane-bound proteins^77^, and altering accessibility of effector interaction surfaces in small GTPases^74,78^.

Interpreting our signaling data (Fig. 5) requires distinguishing the RhoA binding characteristics of the ROCK and mDia effector families, which are two important effector classes of RhoA-mediated actin regulation. For example, the GTPase-deficient RhoA G14V mutant preferentially engages the Rho-binding domain (RBD) of ROCK2 and mDia1 but not that of Rhotekin^79^ while wild type RhoA interacts with all three. This suggests that the G14V GTPase-deficient mutant produces an effector-binding conformation distinct from a GEF-stimulated wild type RhoA.

RhoA-WT-3A increased F-actin and pMLC2 signaling above wild-type levels in both full-serum and serum-starved conditions, whereas RhoA-G14V-3A increased F-actin over RhoA-G14V only in full-serum conditions. A possible interpretation of these data is that G14V saturates the ROCK-pMLC2 axis, so further changes in effector binding competence have little effect, particularly under low stimulation. Structural studies have established that RhoGDI binding to RhoA proceeds through the RhoGDI N-terminal helix-loop-helix domain engaging the switch regions plus α2 and α3 of RhoA, while the RhoGDI C-terminal immunoglobulin-like domain sequesters the C-terminal geranylgeranyl anchor^80^. Analysis of representative GEF, GAP, and effector complexes shows that all major regulatory and effector proteins contact RhoA through the same interface^64^. R5, K6, and K7 are spatially distal from these conserved interaction interfaces, providing structural support for the interpretation that their Ala substitution perturbs the protein-lipid interaction networks rather than regulatory protein binding. Nevertheless, subtle effects on protein-protein interaction cannot be ruled out without direct binding assays.

## Conclusion

In summary, our combined computational and experimental analyses show that RhoA predominantly samples a single membrane organizational state and identify the N-terminal polybasic patch as a secondary membrane anchoring interface. Together with the restricted re-orientation observed here relative to the Ras family proteins, these findings suggest that the structural determinants of membrane orientation dynamics differ across small GTPase subfamilies. Mutation of the N-terminal basic patch increased signaling output without displacing RhoA from the membrane. Instead, it dissolved a cooperative basic-acidic lipid co-binding interaction, released the N-terminus from its dynamically isolated module, and re-poised Switch I toward an effector binding-competent conformation. These results support a model in which the membrane allosterically tunes the conformational competence of the effector lobe.

## Acknowledgements

This work was supported by the National Institute of General Medical Sciences of the National Institutes of Health, grant R01GM144836. Computational resources have been provided by the Texas Advanced Computing Center (TACC) and Anton 2 or Anton 3. Anton computer time was provided by the Pittsburgh Supercomputing Center (PSC) through Grant R01GM116961 from the National Institutes of Health. The Anton 2 and 3 machines at PSC were generously made available by D.E. Shaw Research.

## Author contributions

A.A.G. conceived and designed the project; C.M.H. performed the simulations and GFP-tagged RhoA phalloidin experiments and analyzed simulation data; J.A.F., S.A.P., and Y.Z. performed the HA-tagged RhoA experiments; J.J. performed general cell culture; C.M.H. and A.A.G. analyzed the data and wrote the paper.

