## Supplemental Figures for "Catalytic domain-lipid interactions negatively regulate RhoA signaling by modulating conformational dynamics"

***Analysis of standard structure features.*** Bilayer thickness (P-P) was measured using the z-component of the distance between the centers-of-mass (COM) of the phosphorus atoms in the two leaflets. Area per lipid (APL) was calculated using Voronoi tessellations as implemented in the area-per-lipid class in the Lipyphilic Python toolkit^61^. Probability density functions were calculated using the Gaussian kernel density estimate (KDE) method from the SciPy stats module, with bandwidths assigned using Scott’s Rule. Relative contact frequency is defined as the proportion of frames in which a residue is contacting membrane lipids, where contact is defined as any heavy atom of a residue being within 4.0 Å of a membrane lipid. Hydrogen bonds were measured using an angle cutoff of 135° and a donor-acceptor distance cutoff of 3.0 Å.

To assess RhoA’s membrane orientation, we used two angular reaction coordinates that describe the catalytic domain’s membrane configuration: a tilt angle (θ, 0°-180°), calculated as the angle between the membrane normal and a vector running from the P168 to the D161 Cα atom (V_H5_), and a rotation angle (φ, −180° to 180°) that measures the direction of tilt, defined as follows. First, a reference structure in which V_H5_ was aligned with the membrane normal (N) was generated and a second vector, V_H3_, running from the Cα atom of D161 to the center of mass of the lower portion of Helix 3 (residues 90-91; H_COM_) was defined. We then constructed a rotational reference frame by defining a plane orthogonal to V_H5_ that passes through H_COM_ and projected our reference structure’s V_H3_ onto that plane to obtain V_ref_. We then computed φ as the signed clockwise displacement between the projected V_H3_ and V_ref_ on each frame’s orthogonal plane. This methodology is adapted from one described by Neale et al.^64^ and is commonly used to measure small GTPase orientation^29,65^. 2D histograms of tilt and rotation angles are shown as normalized frequency distributions projected onto a circular plotting surface. To describe small fluctuations in catalytic domain orientation, we quantified the z-distance between the membrane midplane and the COM of the Switch I and α4 regions.

**Control simulation**: While all three residues are solvent-exposed and membrane-facing in the simulations (main Fig 5D-F), K7 was poised to interact with S73 and W58 in some crystal structures (such as PDB ID 1A2B). We therefore ran a separate 10 μs control simulation under conditions identical to those of RhoA-GTP-2 with only R5 and K6 mutated to Ala. The trajectory behaved essentially the same as the triple alanine mutant RhoA-GTP-3A or its wild-type counterpart RhoA-GTP-2. This indicates that mutation of these solvent-exposed residues in tandem did not significantly alter membrane or protein structure or global dynamics. We therefore compared RhoA-GTP-2 with RhoA-GTP-3A in greater detail.

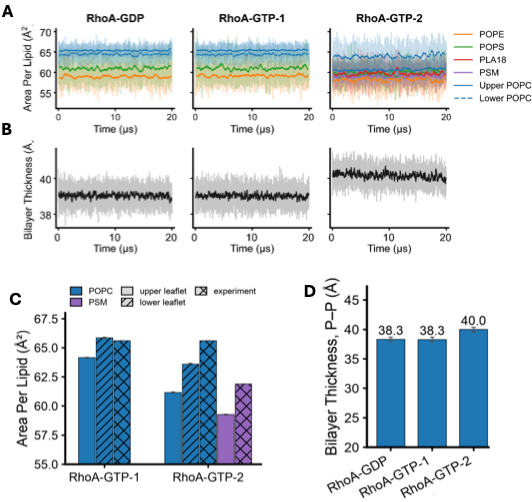

**Figure S1:** Bilayer structural properties. **(A)** Time-evolution of APL for the different lipid species in each simulation. **(B)** Time-evolution of P-P. Heavier lines represent running averages. **(C)** Comparison of ensemble-averaged APL of POPC and PSM from the 3-component (RhoA-GTP-1) and 5-component (RhoA-GTP-2) bilayer simulations with experimental data from pure-component POPC and PSM bilayers. **(D)** Comparison of ensemble-averaged P-P values in the three simulations. APL of POPC in the pure POPC upper leaflet of the asymmetric 3-component bilayer matches the experimental value, validating the simulations. The value for the lower leaflet is slightly smaller, likely reflecting the effect of the PS and PE. The 5-component bilayer behaves in the same manner, with APL of POPC in the upper leaflet being 61.2 Å² versus 63.6 Å² in the PSM-free lower leaflet and 64.2 Å², consistent with the observed increase in membrane thickness. Scattering experiments give pure-bilayer areas per lipid of 65.6 Å² for POPC^1^ and 61.9 Å² for fluid PSM^2^. Ideal, non-condensing mixing of these predicts 63.8 Å² for the 5-component bilayer upper leaflet, whereas the simulated value for the PSM-containing leaflet is ~6% smaller. Sphingomyelin is intrinsically more ordered and more tightly packed than unsaturated phosphatidylcholine^3,4^, consistent with the tighter packing of the saturated-lipid-rich outer leaflet of the plasma membrane^5^. Therefore, the simulation reproduces the increased packing expected when sphingomyelin is present, and is in agreement with our previous simulations of the same membrane composition^6^.

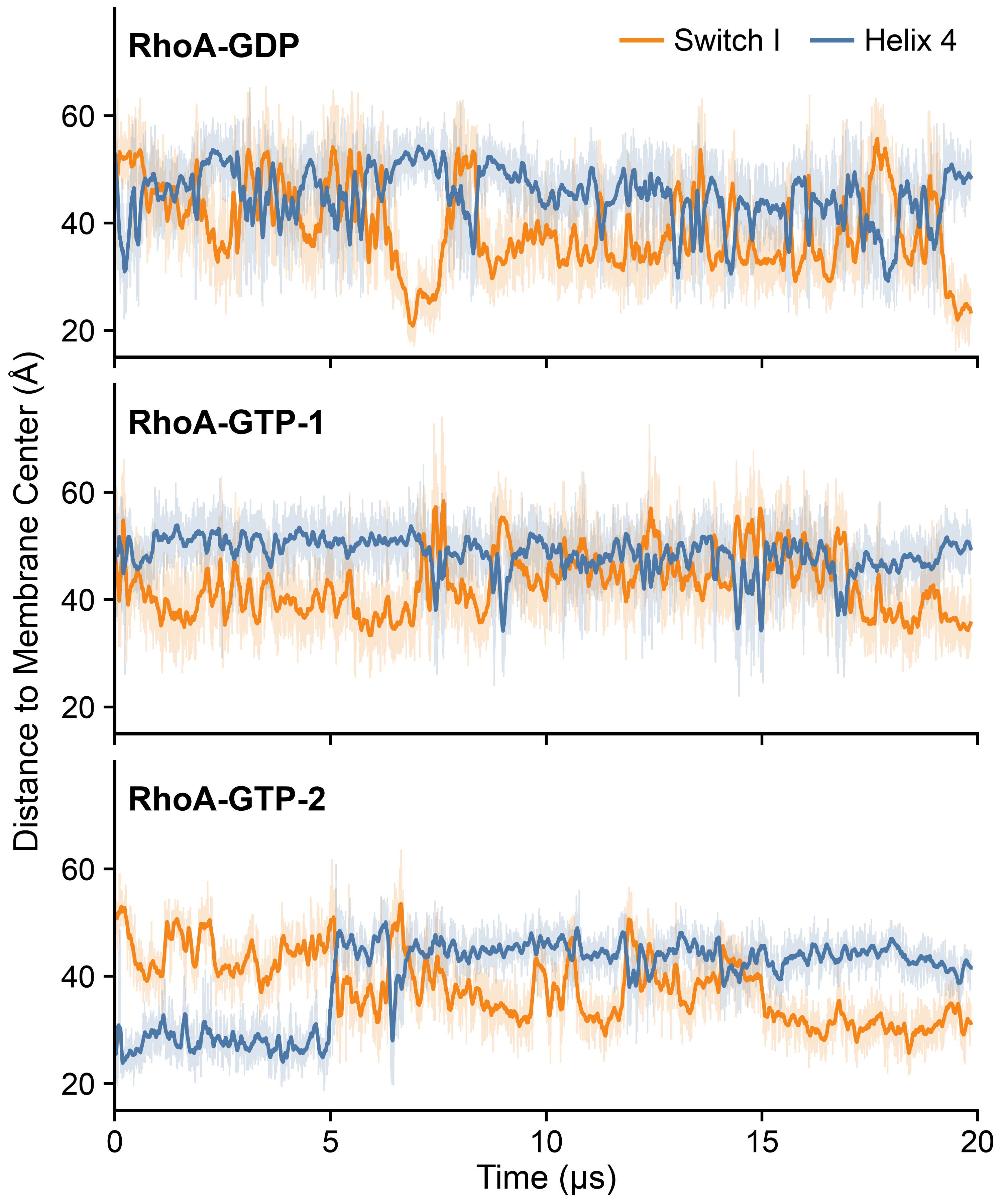

**Figure S2:** Time evolution of the COM Z-distance of Switch I and helix 4 (at opposite faces of the G-domain) from the bilayer center in simulations RhoA-GDP, RhoA-GTP-1 and RhoA-GTP-2.

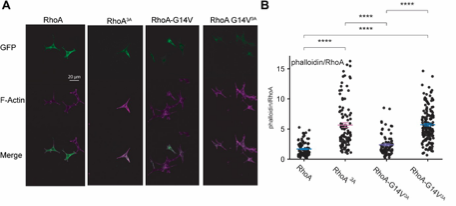

**Figure S3**: Summary of GFP-tagged RhoA F-actin activity. **(A)** Representative confocal microscopy images of NIH 3T3 cells expressing GFP-RhoA and GFP-RhoA alanine mutants under the wild-type and the constitutively active G14V backgrounds. Phalloidin was used to stain for total F-actin (magenta) and GFP expression is shown in green. A 20 µm scale bar is shown in white. **(B)** Quantification of F-actin activity (phalloidin) normalized to GFP-RhoA expression. Each dot represents an individual cell pooled from 3 independent experiments; horizontal bars indicate mean ± SEM. Statistical significance was determined using one-way ANOVA with a post hoc Tukey test. *P < 0.05; **P < 0.01; ***P < 0.001; ****P < 0.0001.
